# A landscape of intergroup conflict shapes den site choice in banded mongooses

**DOI:** 10.1101/2025.09.23.678029

**Authors:** DWE Sankey, T Collins, DP Croft, F Mwanguhya, DW Franks, PA Green, KL Hunt, RA Johnstone, EFR Preston, FJ Thompson, MA Cant

## Abstract

**Background:** For many social animals, intergroup conflict has major impacts on fitness and should therefore influence how groups navigate their environment. Yet most studies of group movement focus on the behaviour of single groups, usually foraging groups studied in isolation of all others. In many systems rival groups present both threats and opportunities, contributing to a landscape of intergroup conflict. This landscape could have a profound impact on group movement.

**Methods:** Here we test how the potential for intergroup conflict influences movement decisions of wild cooperatively breeding banded mongooses, using 14 years of behavioural and GPS data. In this species, encounters between groups can present both risks (e.g. injury/mortality) and opportunities (e.g., extragroup mating). We expected that, to minimise risks and maximise rewards, the motivation to engage in conflict would depend on the group’s state. Such that vulnerable groups (those that were babysitting young offspring) would select den sites closer to the core of their territory than groups without young offspring.

**Results:** We found support for this pattern; non-babysitting groups were unusually risk prone choosing den locations in areas of frequent use by outgroups. During times of heightened recent conflict even choosing to den closer to the core of their rivals’ territory than their own.

**Conclusions:** These results suggest that groups of banded mongooses choose den sites that reflect not only the risks but also the potential rewards of defensive or competitive claims on space. Rival groups thus form an integral part of the social landscape shaping patterns of collective movement.

## BACKGROUND

In humans and many other cooperative vertebrates, encounters between groups result in aggressive intergroup interactions (‘intergroup conflict’) which have major impacts on individual fitness. Intergroup conflict has attracted theoretical attention because of its potential to explain the evolution of altruism within groups^1,2^. However, the influence of intergroup conflict on other aspects of social behaviour, particularly group movement, has been less explored. Groups which lose intergroup conflicts may lose access to vital resources or be displaced into less productive habitats^3,4^. On the other hand, interactions with other groups can bring fitness benefits in terms of opportunities to outbreed^5,6^, win new resources^7,8^, or gain information on reproductive vacancies^9,10^. Other groups thus present both threats and opportunities and can be expected to act as both attractor and repellent influences on group movement^11–13^. The impact of rival groups on movement decisions may also depend crucially on the state of the focal group: groups that are stronger, or stand to gain benefits from intergroup interactions, should be less averse to intergroup encounters or even seek them out, whereas groups that are weak or vulnerable to attack will do best to avoid other groups or areas in which the probability of intergroup interaction is high.

Using 14 years of GPS data from 13 wild banded mongoose (*Mungos mungo*) groups, we tested whether groups adjust a critical spatial decision, their choice of den site, in response to the risks and opportunities posed by rival groups. In this cooperatively breeding species, groups contain multiple females who enter oestrus together and give birth on the same day to a communal litter^14^. Reproduction is synchronised within but not between groups^15,16^. In the first month of life, the communal litter is protected by babysitters who guard the den during the day while the rest of the group goes off to forage^17^. Rival groups pose a deadly risk for babysitting groups because neighbouring groups that find a den containing offspring commonly try to drive off the babysitters and kill the entire litter^18^. Around 20% of pup deaths with a known cause are the result of intergroup infanticide at the den or injuries sustained during intergroup fighting^6^. By contrast, at other stages of the reproductive cycle (i.e. pregnant or non-breeding), intergroup interactions are less common and have less deadly consequences^6^. Thus, we predicted that, when babysitting, groups would try to avoid intergroup interactions (IGIs) by denning in more familiar areas of their territory and in areas that are less commonly frequented by neighbouring groups, compared to when they were not babysitting.

At other phases of the reproductive cycle, intergroup interactions may bring fitness benefits rather than costs, at least for certain group members. Females in oestrus have been observed to initiate intergroup interactions and mate with outgroup males in the midst of battle^5^. Around 18% of pups are fathered to outgroup males in this way. These extra-group pups are heavier, less homozygous, and show better survival in the first year of life than pups fathered by within-group males^6^. Given these potential fitness benefits of intergroup interactions, we predicted that groups containing oestrus females would choose den sites that were more peripheral to their own territory and more core to the territory of a neighbouring group (in terms of frequency of usage), compared to when they were not in oestrus.

In addition, we tested how den-site choice was affected by the frequency of intergroup conflict in the population. At the study site, the frequency of intergroup fighting varies widely between months and years and is particularly common when new groups form and try to carve out a territory that may overlap with the territory of existing groups^19^. In previous studies, the number of IGIs per group per month has varied six-fold from 0.7 to 4.2^16,19^. When the population is stable and food is relatively abundant, groups may go months without any observed intergroup interaction (Cant et al., unpublished data). We used this variation to test whether the effect of babysitting and oestrus on den site choice depended on a group’s recent experience of intergroup conflict in the previous 90 days, which is the approximate length of a reproductive cycle from the start of oestrus to the end of the babysitting period. This biologically meaningful (and data-rich) timeframe allowed us to assess how past conflict might alter risk-sensitive decision-making in den selection.

## METHODS

### Field data

Data were collected over 14 years (Jan 2006-Jan 2020) from a population of wild mongooses found on the Mweya peninsula, Queen Elizabeth National Park, Uganda (0°12’S, 29°54’E). Mongoose groups were visited every 1-3 days and often twice a day throughout the 14-year study period. Group size was noted, as was their location (see section *GPS data* below). Babysitting groups were identified as groups with pups which stayed in the den and did not forage with the group^20^. Often individuals (‘babysitters’) would guard these dens, and the identity of these individuals was also recorded. Oestrus periods were identified through highly conspicuous male mate-guarding behaviour^15,18^. Any intergroup interactions were recorded ad *libitum* (see below: *Measuring outgroup threat*). Rainfall measurements were taken from an on-site weather station.

### GPS data

GPS data were collected from two sources: (1) Handheld GPS Units (2006–2014) – Group locations were recorded every 1–3 days during field observation sessions using Garmin eTrex handheld GPS units; and (2) GPS Collars (2014–2020) – During this period, up to two individuals per group were fitted with GPS collars (weighing 24–41g; Gipsy4 and Gipsy5, Technosmart, Italy).

#### GPS data filtering

GPS data underwent a filtering process to remove erroneous fixes. We excluded collar GPS locations that were associated with less than four satellite connections or with horizontal dilution of precision values (accounting for heteroscedasticity from satellite reception) greater than four^21^. GPS locations were also removed if they fell outside of the feasible range of altitude (<800m or >1100m)^22^.

Where possible, two GPS collars were deployed in each social group to ensure the continuous collection of GPS data in the event of a collar failure. For time periods when two GPS collars were simultaneously collecting data, we only used data from the collar that recorded the greatest number of GPS locations, to avoid problems with duplicated timestamps and locations biasing our territory map estimates^23^. This reduced the number of collar GPS fixes from 83553 to 75992.

In banded mongooses, foraging groups only split when some individuals are left babysitting pups at the den^10,24^. We removed GPS locations from collar data where individuals wearing collars were recorded as babysitters, since this data would not reflect group movement for that period. This decreased the dataset by a further 1017 fixes to 74975.

Finally, all GPS locations (handheld and collar) were overlayed onto a polygon of the shoreline of the Mweya peninsula (e.g., see Fig. 2) and locations that were recorded outside of the boundary of this polygon (i.e. in Lake Edward) were removed. This process reduced the number of GPS fixes from handheld data from 48947 to 47521, and did not remove any fixes from the collar data.

### Den locations

#### Locating dens with GPS

Banded mongoose groups forage during the day and sleep each night in one of many dens in their territory, which are typically abandoned termite mounds or cavities under bushes or rocks. Den locations were recorded using both handheld GPS units and GPS collars (“handheld” N = 478 dens; “collars” N = 326 dens). For GPS data collected using handheld units, the den location was directly confirmed by researchers during the final visit of the day, when the group was observed settling in for the night. For GPS collar data, den locations were inferred based on two consecutive fixes: if the last recorded GPS location of the day (after 1730 hrs) and the first recorded location the following morning (before 0830 hrs) were within 40m of each other, we took the mean of the two as the den location. These time constraints were chosen to align with the stable daylight hours in equatorial Uganda and the natural activity patterns of banded mongooses. The 40m threshold was deemed an appropriate balance between GPS accuracy and the spatial fidelity of den use, as determined in consultation with the field team manager (F. Mwanguhya, pers. comm.).

### Calculating group home-range prior to den event

In R package *ctmm*^25^, we used GPS data to calculate utilisation distributions (UDs), comparable to territories, and the two terms will henceforth be used synonymously. Continuous-time movement models (*ctmm*) calculate kernel density estimates which account for autocorrelation in movement data, i.e., the similarity of data at short time-lags^26^. Models that do not account for autocorrelation have been found to underestimate confidence intervals as well as break the assumption of independence of data points^26–28^. The *ctmm* package produces best-fit models through maximum likelihood chosen by ranked Akaike information criterion (AIC)^25^.

UDs for each mongoose group were calculated from the GPS locations for 90 days prior to the occurrence of each den event, providing a recent picture of the group’s territory up until the day of the located den. The timeframe of 90 days was chosen because it represents one full reproductive cycle for a group (consisting of oestrus, pregnancy and then babysitting^29^). One 90-day cycle is followed by another; there are no significant deviations from this pattern^30^. Therefore, this timeframe i) does not bias toward any particular phase of their reproductive cycle, and ii) is a good estimation of their recent territory use, i.e., larger timeframes may bias towards areas the group previously inhabited but no longer use.

We removed den events where the associated territory had an effective sample size “DOF area”^25^ (where DOF stands for degrees of freedom) lower than 10. This does not indicate 10 GPS data points, but 10 whole crossings of the group’s territory (GPS data required to achieve DOF areas of over 10 are orders of magnitude larger). Special methods are available to deal with DOF areas lower than 10 (see ^31^), and are not required for larger DOF areas, however, we removed den events and the associated territory from our analysis for this data poor category. This removed 44 dens from our analysis (there were 816 dens originally, leaving the 772 den locations we report). We inspected semivariograms to assess the autocorrelation in our data. Semivariograms quantify temporal autocorrelation in movement by plotting the expected squared displacement over increasing time lags, helping to distinguish between range-resident behaviour and unbounded dispersal. In our study, all semivariograms converged rapidly, indicating strong “range-resident” tendencies of banded mongooses. This suggests their movements are spatially restricted, likely due to territory fidelity rather than continuous dispersal^32^.

#### Den locations on a cumulative distribution function (CDF) scale

Having calculated territories for the previous 90 days prior to each den event, we then assessed the dens’ locations in relation to the normal use of the group’s territory. Placing them on a scale of familiarity. For this we use the cumulative distribution function (CDF) of the AIC ranked best-fit UD model and drew the event location from a raster of the CDF data^27,33^(Fig. 2). We further converted CDF to a “familiarity index” by subtracting CDF value from one, as this transformation provided a more intuitive interpretation. After transforming, a familiarity index of 0 means an area infrequently used, and 1 means an area frequently used.

### Outgroup metrics

#### Measuring intergroup interactions

We measured intergroup interactions (defined as any occasion that two groups of mongooses sighted each other and responded by screeching, chasing and/or fighting^16^) as either 1) an occurrence, or not, of at least one IGI over the preceding 90 days, 2) the frequency of IGIs in the preceding 90 days. Intergroup interactions were recorded ad *libitum* during field observations. For each IGI, we recorded the timestamp, the identity of both participating groups and the location coordinates.

#### Measuring outgroup familiarity index

Den sites that are in more familiar/well-used territory do not necessarily confer lower risk of an IGI. For example, where two groups often feed in the same patch, more familiar areas might also come with a higher risk of an IGI. To test whether den choices during babysitting and non-babysitting periods are adjusted to the level of outgroup threat per se, we projected the location of focal group dens onto the territories of rival groups, and calculated an ‘outgroup familiarity index’ over the 90 days before the den event, which measured how familiar a focal group’s den site was to a rival group. Outgroup familiarity index is the CDF of the outgroup’s territory and ranges from 1 in areas of frequent use by outgroup, to 0 for areas of infrequent use. We predicted that groups would choose den sites that are in locations that are more familiar to a rival group when they are non-babysitting (and hence relatively invulnerable to attack) compared to when they are babysitting offspring in the den.

Outgroup familiarity index was calculated for each den for all possible outgroups which existed at the time of the den. If there were two or more outgroups, we took the maximum outgroup familiarity value from any data sufficient territories (DOF area>10). The rationale of using maximum outgroup familiarity and not mean outgroup familiarity, is that if the focal group had several meaningful rivalries during the preceding breeding period, and then denned in a highly familiar area of one of these groups, this effect would be diluted by including the familiarity index of all the other groups territories. The number of IGIs observed over the previous 90 days between the focal group and the (maximum familiarity) outgroup was then calculated, which ranged from no encounters to eight (mean = 0.53).

#### Territory overlap

To calculate the overlap of the focal group’s territory with that of an outgroup, we first found the product of the rasterised *ctmm* models for the focal and outgroup (the group with the highest outgroup familiarity index for the den event) combined. This provided a new overlap model, where the peaks are the areas which both groups use regularly. A grid cell which a first group uses, but a second group rarely uses, will be multiplied by near zero (due to low use by the second group) and will not appear strongly in the overlap territory. We then estimated size of the overlap territory (km^2^, in *ctmm*^25^) and divided the area of the focal group’s home range size by the size of the overlap territory, providing the size of the overlap relative to the size of the focal home range size as a metric of overlap (as a proportion) for analysis.

### Statistical analyses

#### Model 1. Den location on *focal* group familiarity scale against babysitting status

We predicted that groups would respond to the threat posed by outgroups with respect to their babysitting status. Specifically, we expected babysitting (vulnerable) groups to den in familiar territory, while non-babysitting groups would den in less familiar territory.

To assess this, we used den location as the response variable, measured on a familiarity index scale (1-CDF). This was modelled in four separate GLMMs using “glmmTMB”^32^. The models used a beta regression, given the bounded (between 0 and 1) nature of CDF values. We tested five models to compare the effects of babysitting (presence/absence at the time of the den) and IGIs (both as presence/absence and as a frequency over the 90 days prior to the den event) on familiarity index while also considering potential interactions. The models were structured as follows: (1) IGI Presence/Absence + Babysitting (Y/N); (2) IGI Presence/Absence x Babysitting (Y/N); (3) IGI Frequency + Babysitting (Y/N); (4) IGI Frequency x Babysitting (Y/N); (5) IGI Presence/Absence (no babysitting information); (6) IGI Frequency (no babysitting information); (7) Babysitting (Y/N) (no IGI information); (8) Null model with no IGI or Babysitting information. Other fixed effects included in all models were rainfall (mm; mean daily rainfall over the past 90 days), since lower rainfall has been associated with range expansion as mongooses search farther for food^16^; data type: proportion of “handheld” data used to construct the UD; to account for any systematic bias across the two methods; and mean group size over the past 90 days (number of mongooses over 6 months old), which may impact home range size as wider searches may be necessary to find enough food for larger groups. Additionally, all models incorporated a random effect for focal group identity to control for repeated measures and group-specific responses, and an autoregressive correlation structure “ar1()”^34^ where date of the den was nested within group identification ID and babysitting status. This autocorrelation structure reduced autocorrelation in the residuals to beneath the threshold for significance in all models reported in the main text (tested using base-R function “acf()” on the model’s residuals). Analyses were performed in R version 4.3.2^35^.

To determine which model best explained den location on our familiarity index scale, we used an information theoretic approach, using AIC (Akaike Information Criterion) model comparison^36^, where lower AIC values indicate a better trade-off between explanatory power and model complexity. Models were compared using ΔAIC, where values ≤2 indicate similar support, while higher values suggest a weaker fit.

#### Model 2. Den location on *rival* group familiarity scale against babysitting status

We expected babysitting groups to den in familiar territory because we assumed familiar parts of territory were safer from intergroup interactions. However, a more direct way to measure how close dens are to areas of frequent use by rivals is to map denning locations onto rival group territories. This model follows the exact process as model 1, except that the response variable was outgroup familiarity index; fixed effects were: the same four combinations of babysitting (presence/absence) and IGIs (presence/absence) and IGI frequency (but only the frequency of IGIs with the specific outgroup from which we calculated outgroup familiarity index), rainfall, focal group size, and the proportion of “handheld” and “collar” GPS data used to calculate the UD of the rival group. Random effects were the same (focal group identity) because we were interested in controlling for groups which inherently venture more into rival territory than others. Autocorrelation structure, again, was date of the den nested within group identification ID and babysitting status. This autocorrelation structure brought the autocorrelation of the residuals to below significance. Model comparison using AIC scores was run in the same way as Model 1. However, unlike Model 1, the model residuals had a right skew which was corrected for by raising outgroup familiarity index to the power of 1/5.

#### Models 3 and 4: Den location familiarity against oestrus status

Our prediction with regards to oestrus status is that mongoose groups will den in less familiar territory during oestrus periods. This prediction follows the hypothesis that females are motivated to seek outgroup matings and are influential over group movement. To test the impact of oestrus status on den site location on focal group familiar terrain, and rival group familiar terrain, Models 3 and 4 follow Models 1 and 2 (respectively) precisely except that babysitting status (present/absent) was replaced with oestrus status (present/absent).

#### Model 5. Home range estimate

To provide contextual, descriptive statistics regarding factors that increase or decrease home range size, we modelled home range size (km^2^) as a function of IGI (Presence/Absence), rainfall (mm), group size, proportion of data from collars in UD. IGI (Presence/Absence) was used instead of frequency because the AIC comparison revealed it was a better fit. Random effects were group identification. Autocorrelation structure was date of den nested in babysitting status. Home range size was transformed by adding one, applying log, and then raised to the power of 1.6. After transforming, residuals were symmetrical but long tailed. Oestrus (Presence/Absence) and Babysitting (Presence/Absence) were not included as fixed effects because they refer to the group’s state at the time of the den event, and not the territory measured over the previous 90 days.

#### Model 6. Territory overlap

To assess whether more frequent intergroup encounters are associated with greater overlap between focal groups and outgroups, we ran a final model where territory overlap (see methods above) was used as the response variable. This metric is bounded between 0 (no overlap) and 1 (perfect overlap), Thus, we again performed beta regression in a generalized linear mixed model (GLMM)^34^ with a logit link function to appropriately capture the underlying distribution of the data. Here we used all fixed effects used in previous models—IGI frequency (specific to the outgroup whose territory was used in the overlap calculation), babysitting status, oestrus status, rainfall (mm), group size, proportion of collar data in focal UD, proportion of collar data in rival UD—given that these variables were either reasoned to causally influence movement behaviour (which could in turn influence territory overlap) or provided to control for their effects –in the case of proportion of collar data metrics. Random variables included were focal group and rival group identification. Autocorrelation structure was, again, date of the den nested in babysitting status. The model residuals exhibited a right-skewed distribution, which we partially corrected by raising the outgroup familiarity index to the power of 0.7. This transformation improved residual symmetry while maintaining interpretability. Post-transformation diagnostics (Fig. S1) indicated that while the Kolmogorov-Smirnov test still detected some deviation from expected normality, dispersion and outlier tests were non-significant, suggesting no major concerns regarding overdispersion or extreme residual patterns, and residuals were approximately symmetric around zero. Finally, the final statistical test supported the same trend observed in the raw data, as visualised in a basic scatter plot (Fig. S1).

## RESULTS

Home ranges across all measured territories had a mean of 0.251 km^2^ (IQR: 0.119–0.353 km^2^). This was significantly impacted by group size (GLMM; Estimate = 0.0023, S.E. = 0.0008, z = 3.003, p = 0.003) with larger groups holding larger territories. Whether IGIs were present or absent in the previous 90 days, and rainfall, did not have a significant impact on the home range estimate (Table S2).

We found 772 unique denning locations between January 2006 and January 2020, across the 13 mongoose groups we studied (mean dens per group = 55). We found that dens that were located closer together in time were also close together in distance (Fig. 2A). The general pattern was that this was more pronounced in babysitting groups than non-babysitting groups (Fig. 2A). Dens were located throughout each mongoose group’s territory but were concentrated in areas of greater familiarity (Fig. 2B), which could, of course, be more familiar terrain *because of* the presence of dens. Intergroup interactions (IGIs) locations were also distributed across the whole range of a group’s territory. Unlike den sites, IGI locations were concentrated in regions of low familiarity (familiarity index < 0.2; Fig. 1C).

**Figure 1.**
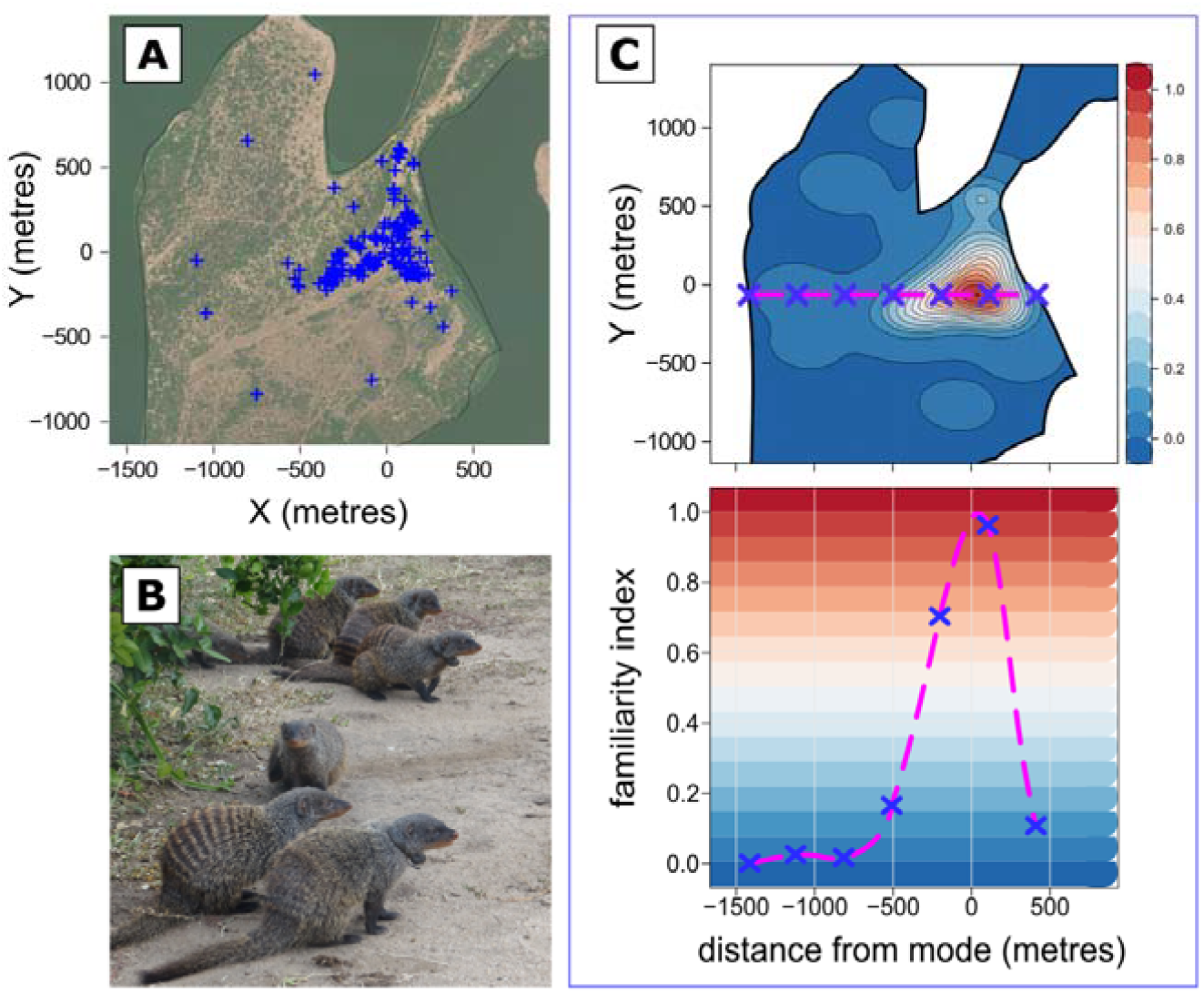
Calculating familiarity index of den sites and IGI locations. **A)** 90 days of GPS data from animal attached collars and handheld GPS units (blue plus signs) for a **B)** banded mongoose group on the Mweya peninsula, Uganda (Photo credit: DWE Sankey). **C)** These data are to calculate a territory in *ctmm*^32^, from which the CDF (cumulative distribution function) can be extracted for each event location of interest (den or IGI). Familiarity index (1-CDF; blue-red scale) describes a focal group’s familiarity of an event location, e.g., den sites or IGI events (blue crosses).

**Figure 2.**
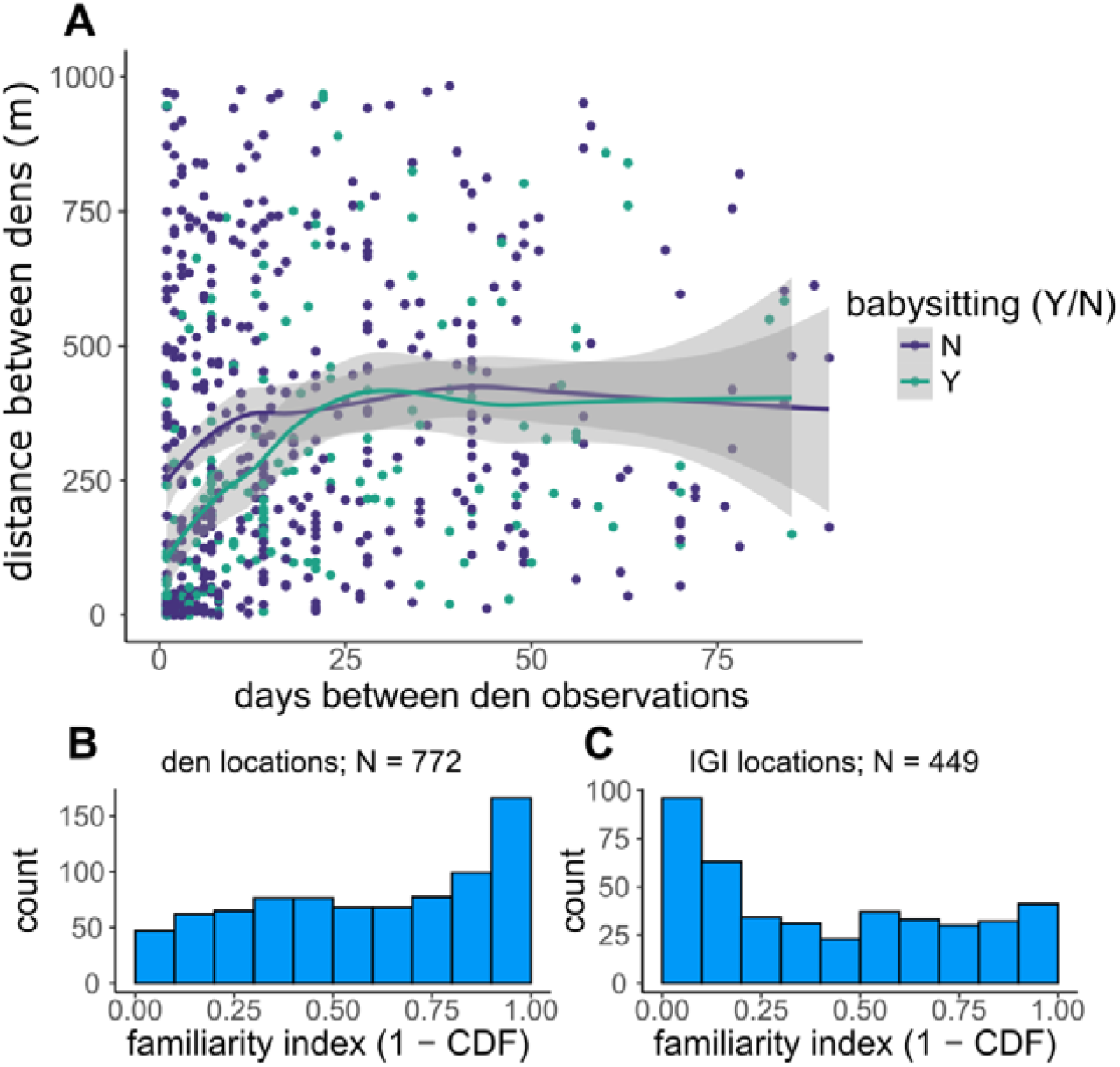
Mongoose den and IGI patterns. **A)** The spatiotemporal autocorrelation of den locations declines with the time between them, as shown by the flattening-off of the relationship between inter-den distance and inter-den time. This decline occurs more slowly in babysitting groups (green) than in non-babysitting groups (purple). Smooth lines are locally weighted smoothed means; grey shaded areas are 0.95 confidence interval^37^. **B-C)** Frequency histograms of Familiarity index (1 – CDF; cumulative distribution function of the estimated territory model) for **B)** Den locations and **C)** Intergroup interaction (IGI) locations.

### Influences on den site (focal group familiarity scale)

Regarding a focal group’s choice of den site relative to their own territory, presence or absence of IGIs over the past 90 days was a stronger predictor of den site familiarity than the frequency of IGIs over the past 90 days (see ΔAIC in Table S1). Specifically, groups den in less familiar territory when there had been a recent intergroup interaction (GLMM: N= 766, Estimate = -0.218, S.E. = 0.1094, z= - 1.9893, p = 0.047; Fig. 3). Babysitting status was also significantly predictive of den site familiarity, with babysitting groups more likely to den in familiar territory (GLMM: N= 766, Estimate = 0.269, S.E.= 0.110, z = 2.444, p = 0.015; Fig. 3). An interaction between babysitting status and IGI (Presence/Absence) was not a more parsimonious fit (see ΔAIC in Table S1), and thus our data supported additive effects of both variables.

**Figure 3.**
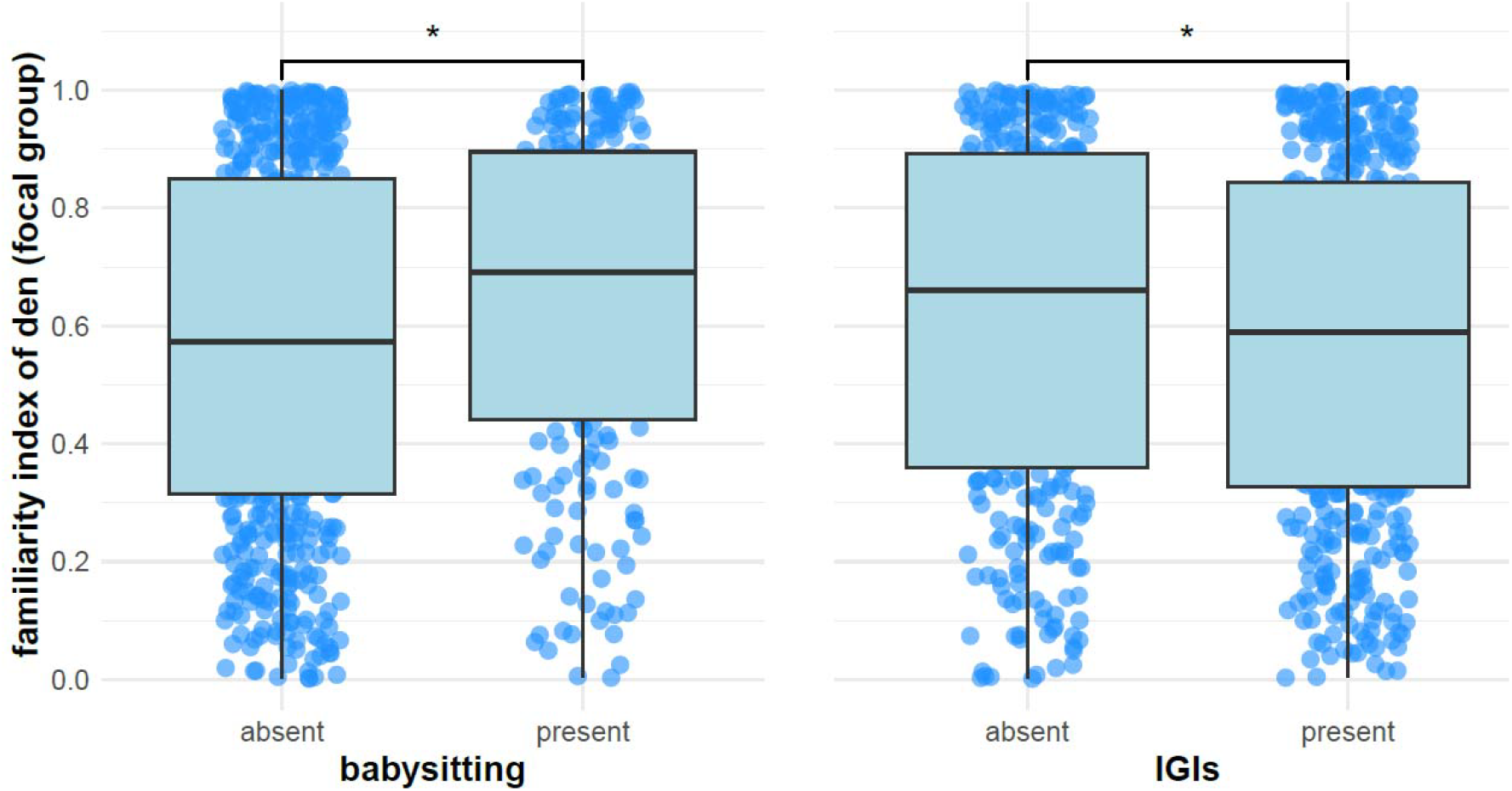
Dens on a scale of focal group familiarity. Boxplots of focal group familiarity index (of a den) over **A)** Babysitting (Presence/Absence) and **B)** Intergroup Interactions (Presence/Absence in the 90 days preceding the den event). The line inside each box is the median den site familiarity, the box represents the interquartile range (IQR), and whiskers extend to values within 1.5 times the IQR. Blue points show individual data. Significance notation (*) represent p values lower than 0.05.

### Influences on den site (outgroup familiarity scale)

We obtained outgroup familiarity scores for 721 of the 772 dens in our dataset. Consistent with our hypothesis that non-babysitting groups choose riskier den sites, we found that non-babysitting groups denned in areas that were more familiar to their rivals compared to babysitting groups (GLMM; N = 716, Estimate = -0.3205, S. E. = 0.1299, z = -2.4674, p = 0.014; Fig. 4A). The frequency of IGIs with the outgroup (for whom the den location had the highest familiarity index score) was a better predictor of outgroup familiarity index than IGI (Presence/Absence) (see ΔAIC in Table S1). Here, higher frequency of IGIs with outgroups was predictive of den site in locations of high familiarity to the same outgroup (GLMM; N = 716, Estimate = 12.378, S.E. = 3.693, z = 3.352, p = 0.001; Fig. 4A). This pattern could partly be due to increase in territory overlap when IGIs were more frequent (GLMM of territory overlap as a function of IGI frequency: N = 721, Estimate = 6.376, S.E. = 2.482, z =2.569, p = 0.010, Fig. S1), which may explain the unexpected result that the dens of vulnerable babysitting groups also show increased outgroup familiarity index during frequent conflicts (Fig. 4A, green line). Overlap alone, however, cannot explain the trend of the non-babysitting groups. When conflicts were most common (> 4 IGIs in the previous 90 days), non-babysitting groups chose to den in locations which were more frequently used by the outgroup than the focal group (Fig 4B, purple line).

**Figure 4.**
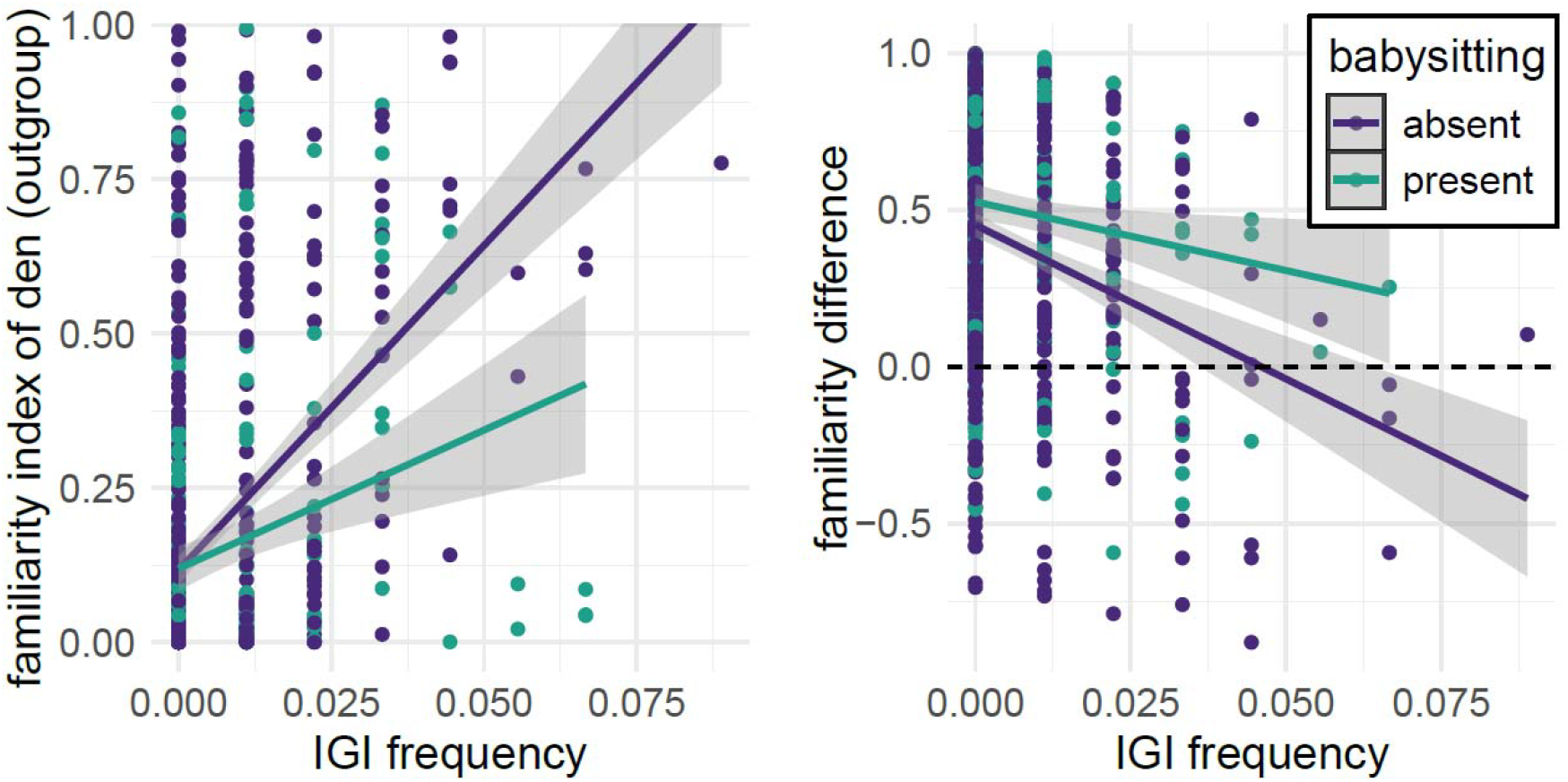
Dens on a scale of outgroup familiarity. **A)** Outgroup familiarity index of den locations against the intergroup interaction (IGI) frequency of the focal and this outgroup during the previous 90 days. The trends – for babysitting groups (green) and non-babysitting groups (purple) – are plotted with ggplot2 “geom_smooth()”, using a linear approximation for each group, which align with the statistical output. **B)** Familiarity difference – the outgroup familiarity index subtracted from the focal group familiarity index of a den – against IGI frequency. Positive values indicate locations more familiar to focal groups, and negative values indicate areas more familiar to outgroups.

### Testing oestrus status influence on den site choice

We found no evidence that groups in oestrus chose den locations in less familiar territory (GLMM: N=766, Estimate = 0.080, S.E. = 0.157, z = 0.510, p = 0.610) or in locations more familiar to the outgroup (GLMM: N=716, Estimate = -0.0831, S.E. = 0.129, z = -0.645, p = 0.519) (Fig. 5). Including (and thus accounting for) babysitting status in this model only weakened the estimated effect of oestrus on familiarity index (focal group familiarity index: N= 766, Estimate = 0.004, S.E. = 0.159, z = 0.030, p = 0.976; outgroup familiarity index: N=716, Estimate = -0.029, S.E. = 0.130, z = -0.221, p = 0.825). Thus, while there is anecdotal and genetic evidence that oestrous females often incite intergroup interactions to obtain outgroup matings^5^, these potential fitness incentives of conflict seeking are not reflected in choice of den site.

**Figure 5.**
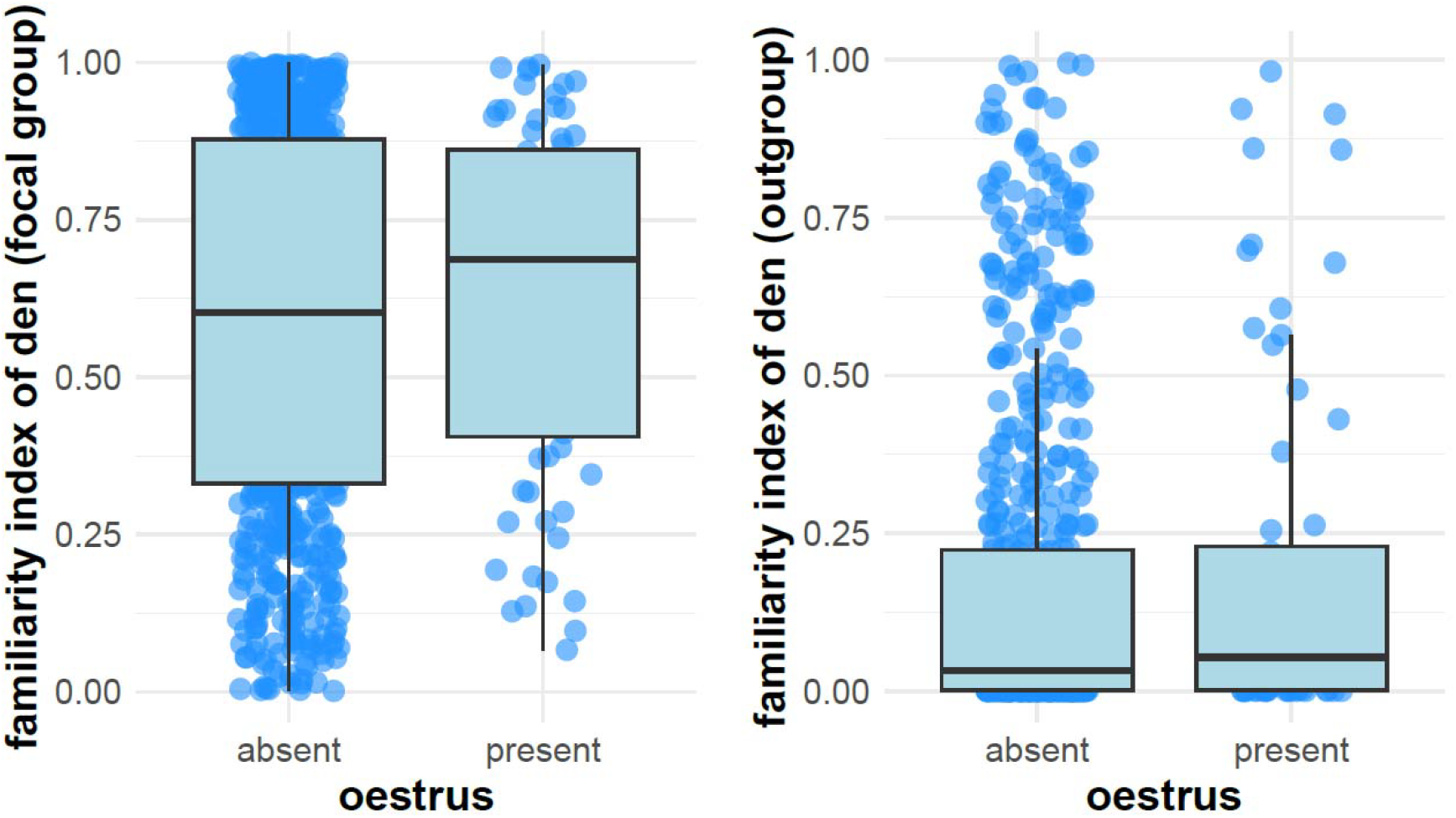
Oestrus state. Our data indicate a lack of evidence that oestrus state impacts den location on a scale of **A)** focal group familiarity and **B)** outgroup familiarity. The line inside each box is the median den site familiarity, the box represents the interquartile range (IQR), and whiskers extend to values within 1.5 times the IQR. Blue points show individual data. Positive values indicate locations more familiar to focal groups, and negative values indicate areas more familiar to focal groups.

## DISCUSSION

To summarise these results, we found that intergroup interactions and babysitting status were predictive of den site choice in banded mongooses. Groups that were babysitting offspring chose den sites that were less frequently used by neighbouring groups compared to when they were not babysitting. As the frequency of past encounters increased, non-babysitting groups chose den sites that were deeper in rival territory (in terms of familiarity). By contrast, we did not find any effect of the group having females in oestrus on den location, suggesting that the potential fitness benefits of outgroup mating did not influence den site choice.

These findings align with our hypothesis banded mongoose groups adjust their choice of den site according to the costs and benefits of intergroup interactions. By denning in areas of high intergroup interaction risk at times when they are particularly strong or invulnerable (for example, when they are not babysitting offspring), banded mongoose groups can gain a wealth of potential benefits, such as the chance to drive off rivals or stake a claim to territory or gain access to new mating opportunities^5^. These findings add to anecdotal reports in banded mongooses that groups sometimes actively seek out conflicts with rival groups^38^. In chimpanzees, intergroup interactions often appear to be the result of active conflict seeking. Male chimpanzees form border patrols which move stealthily into peripheral areas or into a rival group’s home range and attack rival males that they encounter there. Over time males that are successful in killing off outgroup males in intergroup conflict can expand their territory, recruit new females and increase their reproductive success^39^. Similarly, in several species of stingless bees, colonies form ‘fighting swarms’ that seek out and attack the nests of neighbouring groups, to usurp or steal their honey resources^40^. While classic theoretical models of intergroup conflict assume that IGIs occur at random according to some rate parameter^41–45^, our findings add further evidence that that frequency and location of intergroup fights in natural populations reflects strategic choices by groups (or particularly influential individuals within them^46^) to seek or avoid conflict.

It is possible that conflict patterns might be caused by den site choice, rather than den sites being chosen in response to conflict. While correlation does not imply causation, it is unlikely that den site choice is driving conflict patterns in this case. Territories were defined based on the 90 days of movement prior to denning, meaning the spatial context — and any associated conflict — was already established before the den site was chosen. Although the den location might slightly influence future space use, its impact on the preceding 90 days would be minimal, with only the autocorrelation of den sites in space and time (Fig 2A) having influence over the previous (approximately 20) days. Ecologically, it is more plausible that den sites are selected in response to existing territorial dynamics, rather than being the cause of them. Nevertheless, we cannot rule out this possibility, and so in future work, we could strengthen our causal claims by using causal diagrams to guide model structure and by being more deliberate about which variables to condition on, avoiding common pitfalls like adding all predictors into a model or relying solely on AIC-based selection^47^.

Our prediction that groups containing females in oestrus would preferentially den in areas familiar to outgroups was not supported. Although previous findings confirm that females in oestrus have increased motivation to seek outgroup mating opportunities and exert greater influence over group movements^38^, this did not translate into observed patterns of den selection. A possible explanation is that, despite increased female influence during oestrus, conflict-seeking group movements could be constrained by coalitions of males, who may be incentivised to avoid conflict as they incur higher risks from intergroup conflicts^38^. Consequently, male resistance may limit the extent of female-driven movement toward outgroup areas. Future research could investigate whether variation in female-to-male ratios within groups influences directional movement toward or away from outgroup territories, in line with theoretical predictions^48,49^. Additionally, other factors, such as the overriding importance of den-site selection for offspring protection, may have influenced our results.

Our study did not explicitly investigate outcomes of intergroup interactions (i.e., wins or losses) or their long-term consequences. Winning or losing is likely an important factor in governing group space use, given evidence in capuchin monkeys which lost territorial disputes faced restricted access to high-quality food sources^4^, losing baboon groups are displaced from the area of the IGI^50^, and lion prides which lost IGIs were displaced into lower-quality habitats^3^. Future research could extend our findings by explicitly considering the effects of conflict outcomes, investigating how victories or defeats might influence subsequent decisions about den-site selection, territory use, and fitness-related outcomes.

In this study, we used a probability-of-use metric derived from continuous-time movement models (CTMM) to quantify space use by banded mongoose groups. In contrast, other intergroup conflict studies typically quantify space use by examining linear distances or overlaps in ranging areas^3,4^. There is likely to be some general agreement between these two methods, as illustrated in Fig. 1A, since areas of high usage probability often correspond to areas close to the group’s core range. However, key differences also exist. For example, sites like water holes may have high probability-of-use values for multiple groups, yet this overlap might not be accurately reflected by simple distance metrics. A probability-of-use metric offers the advantage of capturing nuanced habitat use patterns across continuous landscapes, whereas distance-based metrics more accurately reflect spatial proximity with less abstraction. Future work might benefit from combining both approaches to fully capture the complexity of intergroup spatial interactions.

Our study underscores the limitations of focusing solely on internal social interactions of a single group as a means to understand group behaviour and collective decision-making^51^. Recent studies have shown the importance of incorporating environmental features alongside internal social interactions to explain patterns of collective decision making^52^. For many social animals, an important and underappreciated component of the social environment is the presence and behaviour of rival groups. The influence of these outgroups is likely to be particularly strong in species, such as social mongooses^30,53^, chimpanzees^39^, wolves^54^, and woodhoopoes^55^, in which interactions between groups involve both substantial risks^5,39^. and opportunities56. landscape of intergroup conflict” in some way. Importantly, these fitness costs and benefits may help to explain when and why conflicts break out, and the extent to which they can be avoided, helping to understand this landscape of intergroup conflict and the dynamics it creates.

## Author contributions

Conceptualization: TC, DWES, KLH, FJT, MAC; Methodology: TC, DWES, FJT, SA, RB, SK, FM, EFRP; Formal analysis: DWES; Resources: MAC; Data Curation: TC, DWES; Writing: DWES, MAC; Visualization TC, DWES; Funding acquisition: MAC, FJT, RAJ, DWF, DPC.

## Consent to Publish declaration

Not applicable

## Ethics

Approval of all work was received from the Uganda Wildlife Authority (UWA) the Uganda National Council for Science and Technology (UNCST), and University of Exeter’s ethical review body.

## Supplementary Material

**Figure S1.**
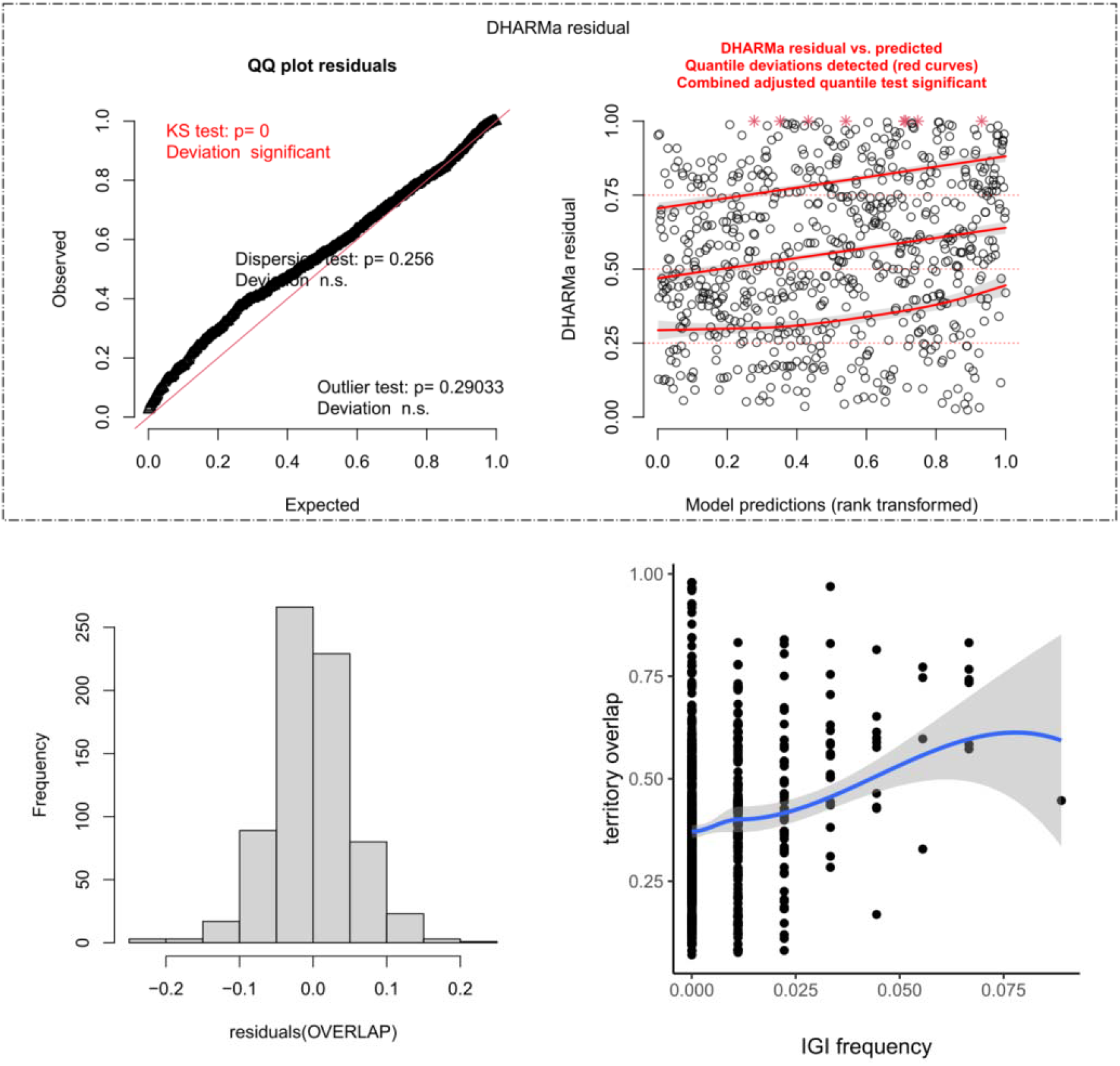
Residual diagnostics and relationship between IGI frequency and territory overlap. (Top left) QQ plot of model residuals, showing a significant deviation from expected normality based on the Kolmogorov-Smirnov (KS) test (p = 0), while dispersion and outlier tests indicate no significant deviations. (Top right) DHARMa^57^ residuals plotted against model predictions, with red curves indicating detected quantile deviations and a significant combined adjusted quantile test, suggesting potential model misspecification. (Bottom left) Histogram of residuals for the overlap model, showing an approximately symmetric distribution centred around zero, despite the overall model exhibiting deviations from expected residual behaviour. (D) Scatter plot of IGI (intergroup interaction) frequency and territory overlap, with a fitted smoothing curve (blue line) from function “geom_smooth()” in ggplot2^58^ using a LOESS method and a grey 95% confidence interval, suggesting a positive relationship with increased variance at higher IGI frequencies. Altogether, despite deviations in residual diagnostics, the histogram of residuals suggests approximate symmetry, and the fitted LOESS curve further supports the trend identified in the statistics, that increased IGI frequency is associated with greater territory overlap (Estimate = 6.376, SE = 2.482, z = 2.569, p = 0.010).

**Table S1.** ΔAIC (Aikaike Information Criterion) scores for different covariate structures for our four hypothesis testing models. Focal and Rival familiarity refers to the familiarity index of the den site in the focal and rival territories respectively. IGI (Y/N) refers to presence or absence of intergroup interactions within the 90 days preceding the den event, whereas IGI (Frequency) is the total number of interactions divided by the 90 days. Babysitting (Y/N) and Oestrus (Y/N) refer to whether the group were babysitting or in oestrus at the time of the den event. D.F. stands for degrees of freedom.

| Model | Structure of Predictor Variables | D.F. | ΔAIC |
| --- | --- | --- | --- |
| **Model 1. Focal familiarity ~ babysitting status** | IGI (Y/N) + Babysitting (Y/N) | 10 | 0.000 |
|  | IGI (Y/N) x Babysitting (Y/N) | 11 | 0.358 |
|  | Babysitting (Y/N) (no IGI information) | 9 | 1.917 |
|  | IGI (Frequency) + Babysitting (Y/N) | 10 | 3.148 |
|  | IGI (Y/N) (no Babysitting information) | 9 | 3.984 |
|  | IGI (Frequency) x Babysitting (Y/N) | 11 | 5.131 |
|  | Null Model | 8 | 5.446 |
|  | IGI (Frequency) (no Babysitting information) | 9 | 6.480 |
| **Model 2. Rival familiarity ~ babysitting status** | IGI (Frequency) + Babysitting (Y/N) | 10 | 0.000 |
|  | IGI (Frequency) (no Babysitting information) | 9 | 4.019 |
|  | Babysitting (Y/N) (no IGI information) | 9 | 9.359 |
|  | IGI (Y/N) (no Babysitting information) | 9 | 9.541 |
|  | IGI (Frequency) x Babysitting (Y/N) | 11 | 0.036 |
|  | IGI (Y/N) + Babysitting (Y/N) | 10 | 1.319 |
|  | IGI (Y/N) x Babysitting (Y/N) | 11 | 3.031 |
|  | Null Model | 8 | 9.099 |
| **Model 3. Focal familiarity ~ oestrus status** | IGI (Y/N) (no Oestrus information) | 9 | 0.000 |
|  | Null Model | 8 | 1.462 |
|  | IGI (Y/N) + Oestrus (Y/N) | 10 | 1.739 |
|  | IGI (Frequency) (no Oestrus information) | 9 | 2.496 |
|  | Oestrus (Y/N) (no IGI information) | 9 | 3.199 |
|  | IGI (Y/N) x Oestrus (Y/N) | 11 | 3.629 |
|  | IGI (Frequency) + Oestrus (Y/N) | 10 | 4.221 |
|  | IGI (Frequency) x Oestrus (Y/N) | 11 | 5.544 |
| **Model 4. Rival familiarity ~ oestrus status** | IGI (Frequency) (no Oestrus information) | 9 | 0.000 |
|  | IGI (Y/N) (no Oestrus information) | 9 | 0.797 |
|  | IGI (Frequency) + Oestrus (Y/N) | 10 | 1.584 |
|  | IGI (Y/N) + Oestrus (Y/N) | 10 | 2.454 |
|  | IGI (Frequency) x Oestrus (Y/N) | 11 | 3.027 |
|  | IGI (Y/N) x Oestrus (Y/N) | 11 | 4.156 |
|  | Null Model | 8 | 4.387 |
|  | Oestrus (Y/N) (no IGI information) | 9 | 5.947 |

**Table S2.**
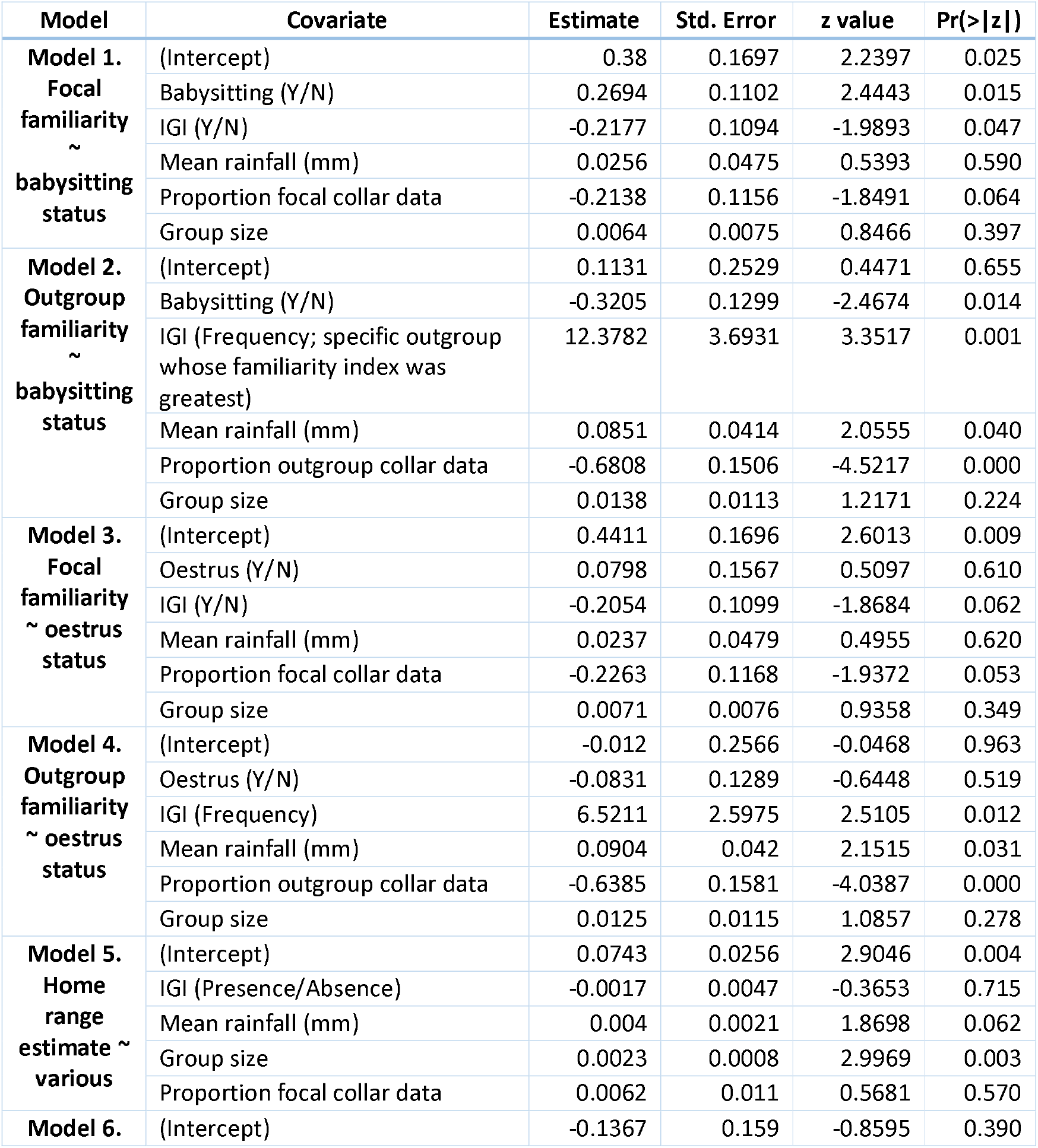
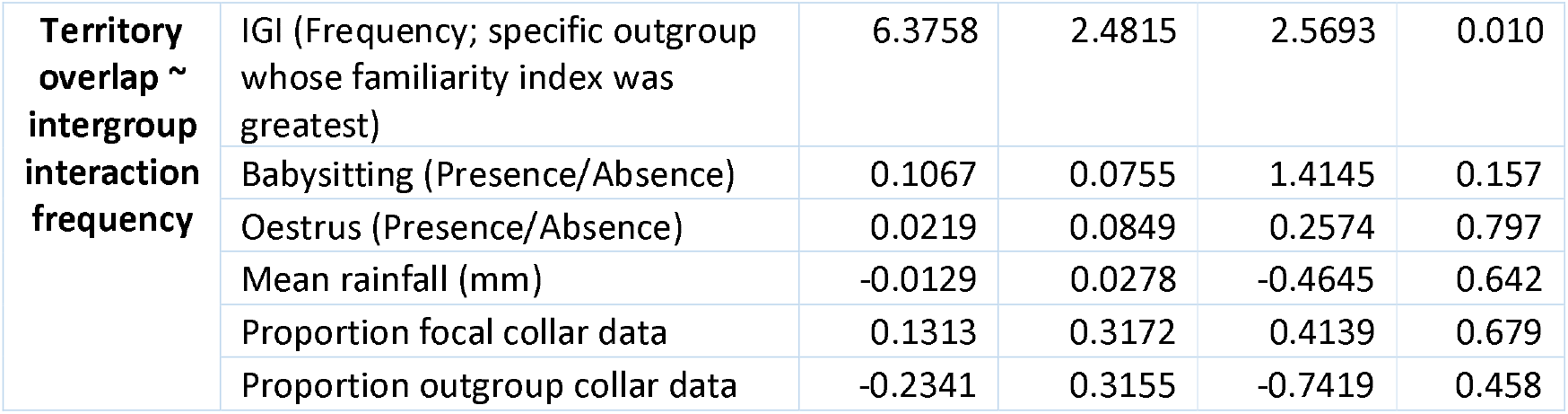
Model statistics from all six models reported in the Main Text. Focal and Rival familiarity refers to the familiarity index of the den site in the focal and rival territories respectively. IGI (Y/N) refers to presence or absence of intergroup interactions within the 90 days preceding the den event, whereas IGI (Frequency) is the total number of interactions in the same time window divided by 90. Babysitting (Y/N) and Oestrus (Y/N) refer to whether the group were babysitting or in oestrus at the time of the den event. Mean rainfall (mm) is mean rainfall over the previous 90 days in mm. Proportion focal collar data refers to the proportion of data used to construct the utilisation distribution (UD) that came from collars, over the total number of GPS points used in the UD, which could be a mixture of “handheld” and “collars” (see Methods). Group size is the number of adults (individuals > 6 months old) in the group. Models are described in the Methods.

